# Fuse to defuse: a self-limiting ribonuclease-ring nuclease fusion for type III CRISPR defence

**DOI:** 10.1101/2020.03.11.987685

**Authors:** Aleksei Samolygo, Januka S. Athukoralage, Shirley Graham, Malcolm F. White

**Affiliations:** Skolkovo Institute of Science and Technology, Bolshoy Boulevard, 30/1, Moscow 121205, Russian Federation; Biomedical Sciences Research Complex, School of Biology, University of St Andrews, North Haugh, St Andrews, Fife KY16 9ST, UK

## Abstract

Type III CRISPR systems synthesise cyclic oligoadenylate (cOA) second messengers in response to viral infection of bacteria and archaea, potentiating an immune response by binding and activating ancillary effector nucleases such as Csx1. As these effectors are not specific for invading nucleic acids, a prolonged activation can result in cell dormancy or death. To avoid this fate, some archaeal species encode a specialised ring nuclease enzyme (Crn1) to degrade cyclic tetra-adenylate (cA_4_) and deactivate the ancillary nucleases. Some archaeal viruses and bacteriophage encode a potent ring nuclease anti-CRISPR, AcrIII-1, to rapidly degrade cA_4_ and neutralise immunity. Homologues of this enzyme (named Crn2) exist in type III CRISPR systems but are uncharacterised. Here we describe an unusual fusion between cA_4_-activated CRISPR ribonuclease (Csx1) and a cA_4_-degrading ring nuclease (Crn2) from *Marinitoga piezophila*. The protein has two binding sites that compete for the cA_4_ ligand, a canonical cA_4_-activated ribonuclease activity in the Csx1 domain and a potent cA_4_ ring nuclease activity in the C-terminal Crn2 domain. The activities of the two constituent enzymes in the fusion protein cooperate to ensure a robust but time-limited cOA-activated ribonuclease activity that is finely tuned to cA_4_ levels as a second messenger of infection.

## Introduction

Type III CRISPR systems have class 1 effector complexes that utilize CRISPR RNA (crRNA) to detect invading RNA from mobile genetic elements (MGE) such as viruses. This target RNA binding results in the activation of the Cas10 subunit, which commonly harbours two active sites: an HD nuclease domain for ssDNA cleavage (1-3) and a PALM polymerase domain that cyclises ATP to generate cyclic oligoadenylate (cOA) molecules (4-7). cOA acts as a second messenger in the cell, signalling infection and activating a range of auxiliary defence enzymes such as the ribonuclease Csx1/Csm6 (8-10) or the DNA nickase Can1 (11) by binding to a CRISPR-associated Rossman fold (CARF) sensing domain. Once activated, these enzymes degrade both host and invading nucleic acids in the cell, which can result in viral clearance (i.e. immunity) or lead to cell dormancy or even death (akin to Abortive Infection) (12). Detection of 1 viral RNA by the *Sulfolobus solfataricus* type III-D CRISPR system can lead to the generation of 1000 molecules of cyclic tetra-adenylate (cA_4_), equivalent to an intracellular concentration of 6 µM cA_4_, which is sufficient to fully activate the Csx1 ribonuclease (13).

The potency and high amplification factor inherent in type III cA_4_ signalling necessitates a mechanism to clear cA_4_ from the cell once a viral infection has been defeated – otherwise cells could be driven to “commit suicide” unnecessarily. In *S. solfataricus*, cells express a specialised catalytic variant of a CARF-domain protein named “CRISPR-associated ring nuclease 1” (Crn1) that binds and slowly degrades cA_4_, returning cells to a basal, uninfected state (14). The CARF domains of some members of the Csm6 family have also been shown to have ring nuclease activity, slowly degrading cA_4_ and thus acting as self-limiting ribonucleases (9,15). More recently, a potent ring nuclease enzyme of viral origin has been identified that acts as an anti-CRISPR (Acr) by rapidly degrading cA_4_ to circumvent type III CRISPR immunity (16). The enzyme, AcrIII-1, is a small dimeric protein unrelated to the CARF domain that uses an active site histidine residue to ensure higher catalytic efficiency (16). Biochemical and modelling studies have revealed that AcrIII-1, which is widespread in archaeal viruses and bacteriophage, efficiently deactivates Csx1 *in vitro* (17).

Homologues of AcrIII-1 are also found in association with type III CRISPR systems in some genomes, where they have been predicted to function in the host defence against MGE. In this context, the enzyme has been named Crn2 (CRISPR-associated ring nuclease 2) (16). We previously noted an unusual example of a Crn2 gene fused to a Csx1 ribonuclease in the type III CRISPR locus of the organism *Marinitoga piezophila* (16), and herewith we term this gene product Csx1-Crn2 (Figure 1). The *M. piezophila* CRISPR locus includes a type III-B effector, an RNA-guided Argonaute enzyme (18) and an uncharacterised CARF-RelE protein that may be activated by cOA to cleave ribosomal A-site mRNA (19). Sequence analysis suggests that Csx1-Crn2 consists of a canonical Csx1 protein, including an N-terminal CARF domain for cOA sensing and HEPN (Higher Eukaryotes, nucleotide sensing) ribonuclease domain, fused to a Crn2 domain at the C-terminus (Figure 1B). The 563 aa protein is predicted to form a dimer, and a structural model generated by Swissmodel (17) using the templates Csx1 (PDB 6R7B) and AcrIII-1 (PDB 6SCF) is shown in Figure 1C.

**Figure 1.**
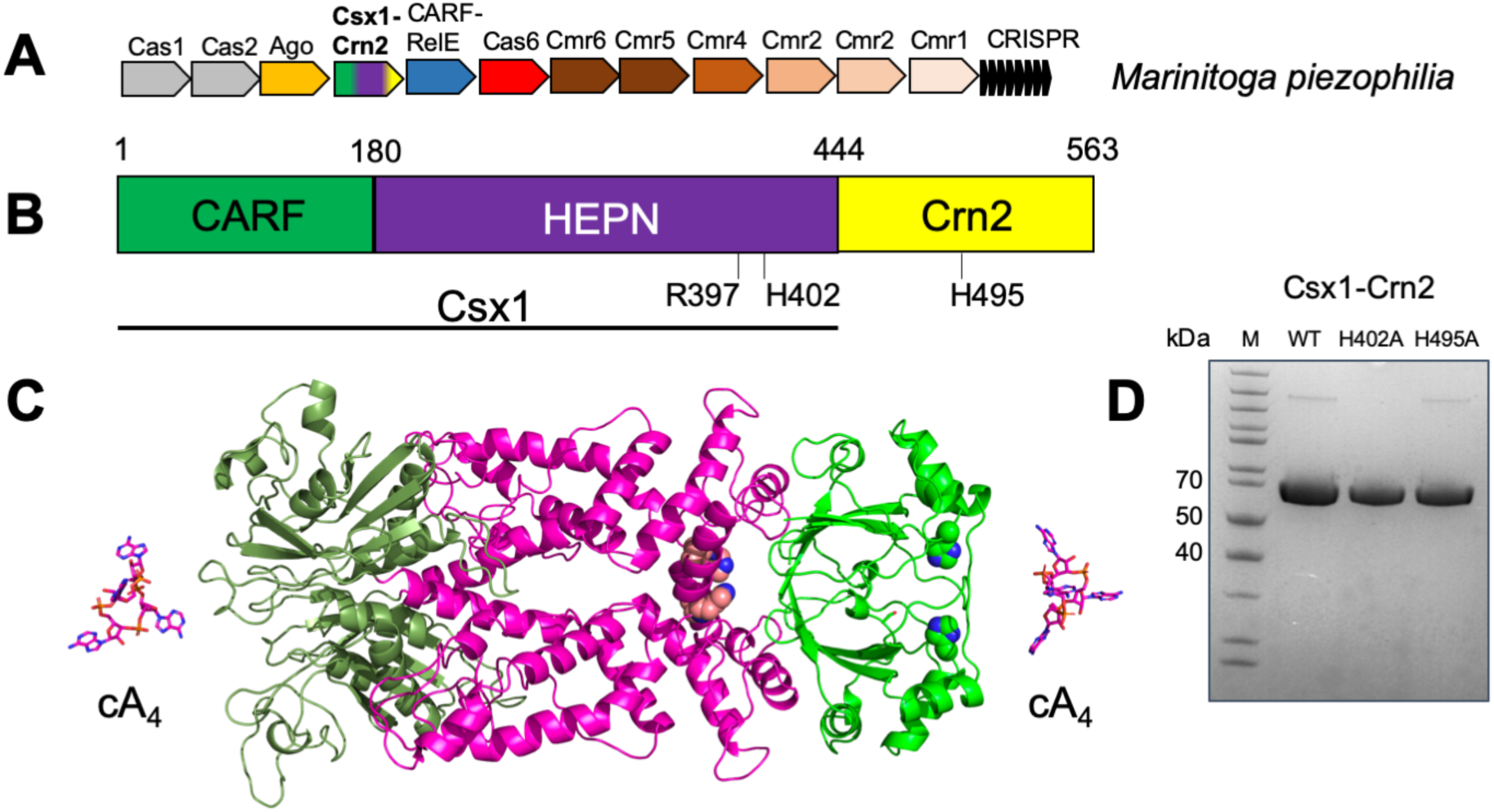
The atypical type III CRISPR system of M. piezophila. **A.** CRISPR locus of M. piezophila includes genes encoding a complete type III-B (Cmr1-6, Cas6) gene cluster, adaptation proteins Cas1 and Cas2, argonaute, and two uncharacterised CARF-domain proteins: CARF-RelE and Csx1-Crn2. **B.** Cartoon of the Csx1-Crn2 protein highlighting the N-terminal Csx1 ribonuclease and C-terminal Crn2 ring nuclease components. **C.** Structural model of Csx1-Crn2. The active site residues of the HEPN ribonuclease (R397, H402) and ring nuclease (H495) are shown in spheres. Domain colouring is as for part B. **D.** SDS-PAGE analysis of purified recombinant Csx1-Crn2 wild-type and variants.

Here, we express, purify and characterise the Csx1-Crn2 protein, revealing cA_4_-activated ribonuclease activity associated with the Csx1 domain and a potent cA_4_ ring nuclease activity catalysed by the Crn2 domain – the first cellular ring nuclease of this family to be studied. Biochemical analysis of the wild-type and engineered variant forms of the enzyme allow the mechanism of this self-limiting antiviral defence enzyme to be delineated.

## RESULTS

### Expression and purification of Csx1-Crn2

A synthetic gene (g-block) encoding the *M. piezophila* Csx1-Crn2 protein, codon optimised for expression in *Escherichia coli*, was cloned into the pEV5hisTev plasmid (20), allowing expression of the recombinant protein with a cleavable N-terminal polyhistidine tag. The protein was purified to near homogeneity by a combination of immobilized metal affinity and size exclusion chromatography (Figure 1D). Enzyme variants with mutations targeted to the active site of the Csx1 ribonuclease (H402A) or the Crn2 enzyme (H495A) were expressed and purified as for the wild-type enzyme.

### The N-terminal domain of Csx1-Crn2 is a cA_4_-activated ribonuclease

Type III CRISPR systems typically synthesis a range of cOA molecules with a ring size varying from 3-6 (4-7,21). We first wished to determine the identity of the activator for Csx1-Crn2 by incubating the enzyme with an RNA substrate (RNA A1) in the presence of cA_3_, cA_4_ or cA_6_. Only cA_4_ activated the ribonuclease activity of the enzyme (Figure 2A), consistent with the predicted structural similarity to the cA_4_-activated Csx1 protein of *S. islandicus* (10). A variant enzyme with a mutation targeted to the HEPN ribonuclease active site (H402A) was inactive for RNA degradation, but the H495A variant with a mutation targeted to the Crn2 active site appeared more active than the wild-type enzyme (Figure 2B). Overall, these data confirmed that the HEPN domain of Csx1-Crn2 was responsible for RNA cleavage and suggested that the ring nuclease activity might serve to limit the ribonuclease activity.

**Figure 2.**
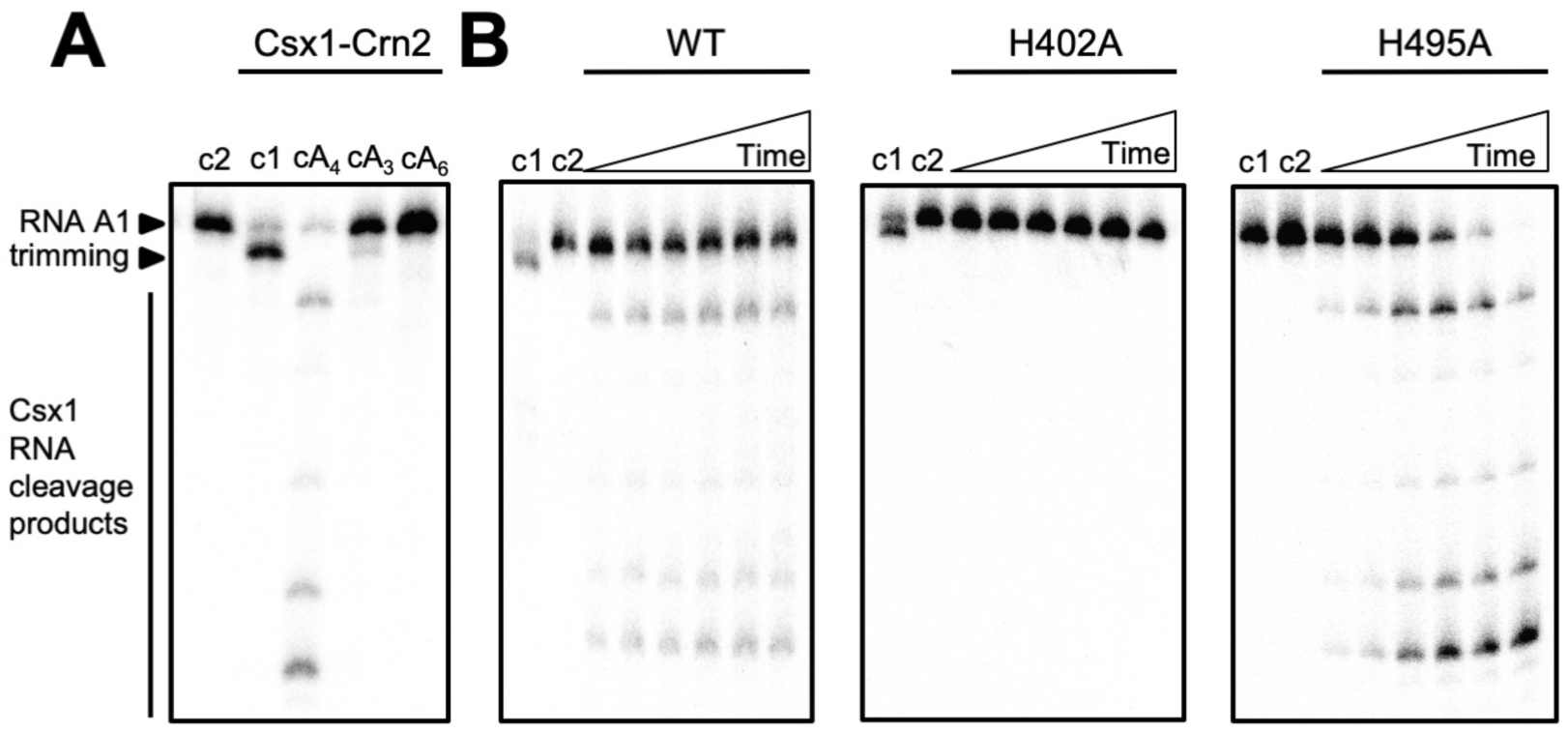
The Csx1 domain of Csx1-Crn2 is a cA_4_-activated ribonuclease. **A.** RNase assay to determine activator of Csx1 domain. Phosphorimage of denaturing PAGE shows the radiolabeled RNA A1 assayed with 1 µM Csx1-Crn2 dimer and a cyclic activator (100 µM): cA_4_ (cyclic tetra-adenylate), cA_3_ (cyclic tri-adenylate), or cA_6_ (cyclic hexa-adenylate). Samples were incubated at 50 °C for 30 min. The control lanes c1 and c2 show the assay in the absence of activator or the absence of enzyme, respectively. The data is representative of three technical replicates. **B.** Phosphorimage of denaturing PAGE visualising degradation of radiolabeled RNA by Csx1-Crn2 or its variant H402A (10 µM dimer) when activated by cA_4_ (100 µM) at 50 °C (time points are 1, 2, 4, 10, 15, 30 min). Control reactions carried out to the end-point of each assay without cA_4_ (c1) or without enzyme (c2) are shown. The data is representive of three technical replicates.

We noted that the RNA substrate was processed in the absence of cA_4_ activator by trimming of a small number of nucleotides from the 3’ end (e.g. Figure 2A lane 2). This activity was abolished in the H495A variant (e.g. Figure 2B lane c1) suggesting that the ring nuclease domain is capable of processing the 3’ ends of RNAs independent of cA_4_ activation. This RNase activity is reminiscent of the CARF-family protein Csx3, which has been described as a deadenylase (22) and has parallels with the observed binding of *S. islandicus* Csx1 to the 3’ tetra-adenylate tails of mRNA (23). The 3’-terminus of the RNA A1 substrate consists of four purine nucleotides (GAGA).

### The C-terminal domain of Csx1-Crn2 is a highly active ring nuclease specific for cA_4_ degradation

Having established that cA_4_ was the activator of the Csx1 domain, we tested the ability of the enzyme to degrade cA_4_ *in vitro* (Figure 3A). Rapid degradation of cA_4_ to linear A_2_>P (di-adenylate with a 3’-cyclic phosphate) was observed within the first minute of the assay, followed by a slower conversion of A_2_>P to A_2_-P (di-adenylate with a 3’-phosphate). Overall, the reaction rate and products were highly reminiscent of the AcrIII-1 enzymes studied previously (16). A variant enzyme with a mutation targeted to the predicted active site histidine of the Crn2 domain (H495A) did not degrade cA_4_, confirming the Crn2 domain as the site for ring nuclease activity. The single-turnover reaction kinetics of the Csx1-Crn2 ring nuclease activity were too rapid to quantify, suggesting a catalytic rate constant of the order of 10 min^-1^. To explore this more fully, we carried out a series of kinetic experiments under multiple turnover conditions, using 256 µM cA_4_ substrate and varying the concentration of Csx1-Crn2 from 0.16 nM to 5.12 µM (Figure 3B). The initial reaction rate at each enzyme concentration was quantified and plotted, yielding data that could be fitted with a linear fit of slope 8.8 ± 0.3 min^-1^ (Figure 3B). Assuming that the enzyme is saturated at 256 µM cA_4_, this affords an estimation of k_cat_ for the ring nuclease activity of about 9 min^-1^ – in good agreement with the single turnover data for Csx1-Crn2 and other related ring nucleases (16). Thus, the Crn2 active site is a highly active ring nuclease capable of rapid, multiple-turnover degradation of the cA_4_ activator and does not appear to be significantly rate-limited by substrate binding or product release.

**Figure 3.**
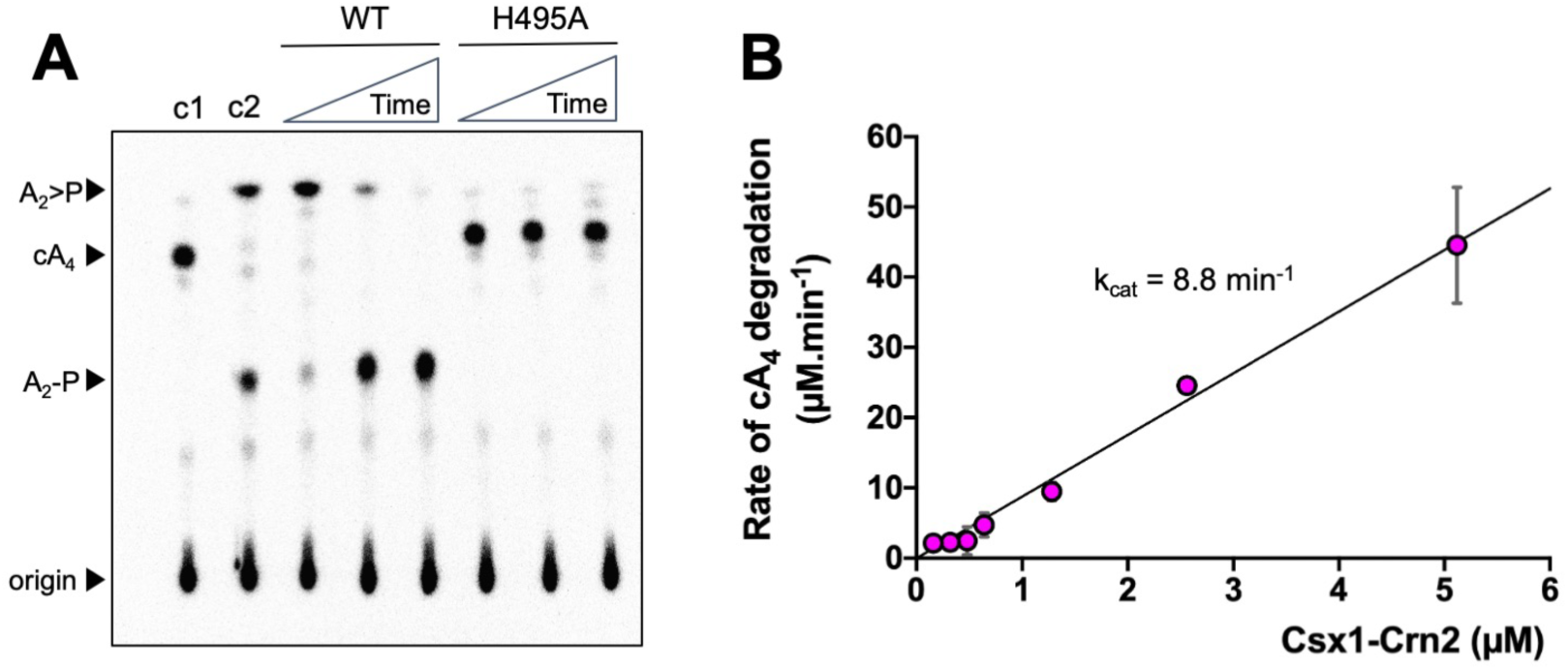
Csx1-Crn2 rapidly degrades cA_4_. **A.** Phosphorimage of thin-layer chromatography (TLC) visualising degradation of radiolabelled cA_4 (_∼200 nM) by Csx1-Crn2 or its H495A variant (1 μM dimer) at 50 °C over time (1, 10, 60 min). Control reactions incubating radiolabelled cA_4_ with buffer (c1, negative control) for 60 min or with 1 µM AcrIII-1 for 1 min (c2, positive control) are also shown; the data are representative of three technical replicates. **B.** Csx1-Crn2 rapidly degrades cA_4_ under multiple-turnover conditions. The initial rate of cA_4_ (256 µM) degradation was quantified at different concentrations of Csx1-Crn2 protein and plotted. A linear function was fitted through the origin and the catalytic rate constant k_cat_ was estimated (k_cat_ = 8.8 ± 0.3 min^-1^). Each data point is the average of at least three technical replicates and error bars show standard deviation of the mean.

### Deactivation of the ring nuclease activity enhances ribonuclease activity of Csx1-Crn2

Having established the activities of the Csx1 and Crn2 domains individually, we next wished to investigate how the two enzymatic domains collaborate to potentiate cA_4_-activated RNA degradation. Firstly, we pre-incubated the wild-type or H495A variant forms of the enzyme with cA_4_ for 10 min, then added wild-type enzyme together with radiolabelled substrate RNA to assess Csx1 activity (Figure 4A). When the (ring nuclease inactive) H495A variant was used in the pre-incubation, robust RNA degradation by Csx1 was observed at all three cA_4_ concentrations tested (Figure 4B). However, following pre-incubation with wild-type Csx1-Crn2, RNA degradation was minimal except for the highest concentration of cA_4_ (100 µM). This experiment demonstrated that Csx1-Crn2 can auto-deactivate by degrading the cA_4_ activator in the ring nuclease site.

**Figure 4.**
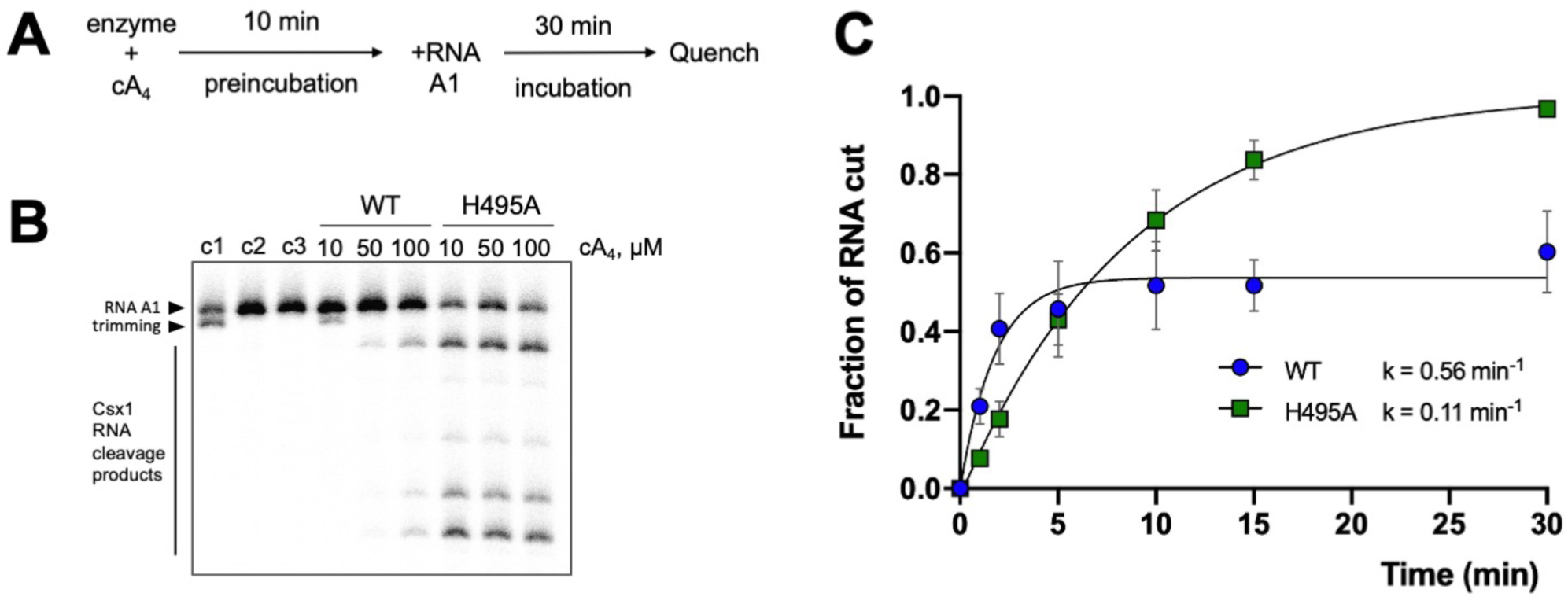
Csx1-Crn2 is an example of a self-limiting ribonuclease. **A.** The scheme of the pre-incubation experiment. **B.** Denaturing PAGE shows degradation of radiolabelled RNA by WT or its H495A variant (10 µM dimer) that were pre-incubated with 10, 50, or 100 µM cA_4_ at 50 °C for 10 min prior addition of radiolabelled RNA A1, followed by incubation for a further 30 min. A set of control reactions in the same conditions but lacking cA_4_ (c1), enzyme (c2) or both (c3) are shown. The data represent the results of three technical replicates. **C.** Quantification of fraction of RNA degraded by either Csx1-Crn2 (WT) or its H495A variant (10 µM dimer) at 50 °C over time. The results were fitted with an exponential equation and estimated rates of RNA cleavage are shown (WT, k = 0.56 ± 0.11 min^- 1^, plateau = 0.54 ± 0.03; H495A, 0.11 ± 0.01 min^-1^, plateau = 1.02 ± 0.03). The data points are the average of three technical replicates and error bars show standard deviation of the mean.

We next quantified RNA degradation over time in a reaction including 10 µM Csx1-Crn2 (wild-type or H495A variant), 100 µM cA_4_ and ∼50 nM radioactively labelled substrate RNA. The wild-type enzyme displayed more rapid initial kinetics, degrading approximately 50% of the RNA within 2 min, but quickly reached a plateau after which RNA was not further degraded (Figure 4C). In contrast, the H495A variant, which is unable to degrade cA_4_, had a slower rate constant but completely degraded all the substrate RNA in the course of the 30 min incubation. Thus, the two active sites have the ability to cooperate to ensure that cA_4_-stimulated RNA degradation is regulated to limit the cellular response to viral infection.

### cA_4_ binding affinity of the CARF and Crn2 domains

Next, we wished to understand the interaction between cA_4_ and the CARF and Crn2 domains of Csx1-Crn2 that bind the second messenger. We first investigated cA_4_ binding by the H495A variant, which can bind but not cleave cA_4_ in the Crn2 site, by electrophoretic mobility shift assay (EMSA), using ∼10 nM radioactively-labelled cA_4_ (Figure 5B). A clear gel-shift was observed yielding an apparent K_D_ of around 20 nM, which is similar to that previously observed for stand-alone ring nucleases (13). As there are two cA_4_ binding sites in the protein, we next designed variant enzymes to abolish cA_4_ binding in either the CARF (H129W) or Crn2 (S452Y) binding sites, guided by the structural model shown in figure 1. These bulky amino acid substitutions were predicted to prevent or significantly reduce cA_4_ binding without destabilising the protein structure. The two variants were purified as for the wild-type protein, and cA_4_ binding was assessed by EMSA (Figure 5B). The S452Y variant showed no obvious binding of cA_4_, but cleavage of cA_4_ at high protein concentrations (apparent K_D_ ∼ 1 µM) suggested that the cA_4_ binding at the ring nuclease site was strongly reduced but not abolished. The H129W variant cleaved cA_4_ into A_2_>P at a concentration of 20 nM enzyme, consistent with high affinity binding and cleavage at the Crn2 site. Overall, these data suggest that the CARF domain has a much weaker intrinsic affinity for cA_4_ than the ring nuclease domain. Otherwise, we would have expected to see higher affinity binding of cA_4_ by the CARF domain of the S452Y variant.

**Figure 5.**
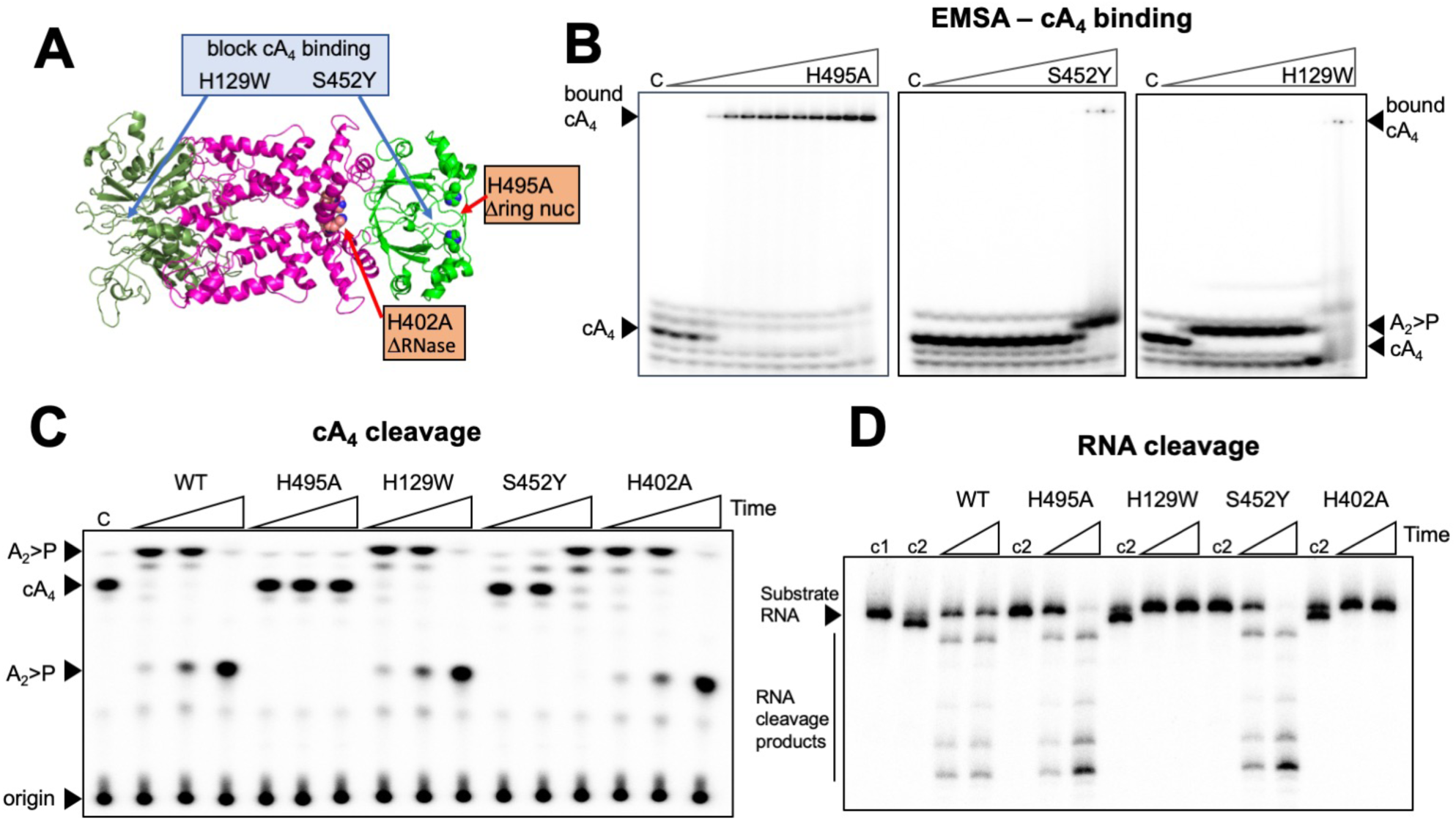
Dissection of cA_4_ binding and catalysis by Csx1-Crn2. **A.** Cartoon representation of Csx1-Crn2 showing sites targeted by mutagenesis. **B.** Electrophoretic mobility shift assay (EMSA) showing binding of radiolabelled cA_4_ (∼10 nM) by Csx1-Crn2 variants (0.001, 0.01, 0.02, 0.03, 0.04, 0.05, 0.06, 0.08, 0.1, 1.0, 10.0 and 20.0 µM, respectively). On each gel is a control reaction (c) with cA_4_ in the absence of protein. The ring nuclease defective variant H495A bound cA_4_ with an apparent dissociation constant around 20 nM. The S452Y variant, which was designed to abolish cA_4_ binding by the ring nuclease domain, had a severe binding defect, with cA_4_ degradation apparent at high protein concentrations. The H129W variant, designed to abolish cA_4_ binding in the CARF domain, showed cA_4_ degradation starting from around 20 nM enzyme. The data are representative of three technical replicates. **C.** TLC showing degradation of cA_4_ (200 nM) by wild-type and variant enzymes (5 µM dimer) at 50 °C. A time-course of 10 s, 1 min and 10 min was carried out and a control reaction (c) incubating buffer with radiolabelled cA_4_ for 10 min is also shown. The H495A variant was inactive as shown previously. The H129W and H402A variants targeting the CARF and HEPN (Csx1) domains, respectively, had normal ring nuclease activity. Whereas the S452Y variant targeting the Crn2 domain had strongly reduced activity, consistent with weaker binding of cA_4_ by the Crn2 domain. **D**. Denaturing PAGE showing cleavage of radiolabelled substrate RNA (100 nM) by Csx1-Crn2 and variants (5 µM dimer) at 50 °C. RNA cleavage was examined at two time-points (5 & 30 min) and control reactions incubating radiolabelled RNA in buffer (c1) or RNA and protein in the absence of cA_4_ activator (c2) for 30 min are also shown. The H129W and H402A variants were inactive as expected. The data are representative of three technical replicates.

We also tested the ring nuclease and RNA cleavage activities of the two variant enzymes (Figure 5C,D). As predicted, the S452Y variant had a severely reduced, although not abolished, ring nuclease activity – consistent with the EMSA result. The H129W variant was a fully active ring nuclease, ruling out a model whereby the two cA_4_ binding sites communicate to control cA_4_ degradation. As expected, the H129W variant had no cA_4_-activated RNase activity, confirming that this variant does not bind cA_4_ in the CARF domain. Neither of the variants targeting the ring nuclease domain had a discernible effect on the activity of the Csx1 ribonuclease.

## DISCUSSION

The importance of cyclic nucleotide signalling in prokaryote anti-viral defence systems is an emerging paradigm. Examples include cA_4_ and cA_6_ for type III CRISPR (5,6), cA_3_ for the HORMA-DncV-NucC system (24,25) and a variety of cyclic dinucleotides synthesised by diverse CBASS enzymes (26,27). Viruses have a clear imperative to degrade these molecules and circumvent immunity, but it is increasing apparent that cellular immune systems also need a mechanism to turnover these second messengers if they are to prevent runaway toxicity and cell death (13). For type III CRISPR systems at least, immunity rather than abortive infection may be the *modus operandi*, as mechanisms exist to “kill the messenger”. Ring nucleases that degrade cA_4_ include the dedicated enzymes and self-limiting ribonucleases based on the CARF domain (9,14,15) and the DUF1874 family whose members include the viral anti-CRISPR AcrIII-1 and putative cellular Crn2 proteins (16).

Crn2 ring nucleases have been detected in diverse bacterial genomes associated with type III CRISPR defence but not yet characterised. Here we focussed on an unusual fusion of a Csx1 ribonuclease and Crn2 ring nuclease from *Marinitoga piezophila*, whose genome also has a CRISPR-associated Argonaute enzyme and an uncharacterised, predicted cA_4_-activated RelE toxin (Figure 1). We have demonstrated that the fused protein does indeed combine the activities of a canonical cA_4_-activated Csx1 ribonuclease and a highly active ring nuclease. The ring nuclease activity, with a k_cat_ of 9 min^-1^, is on a par with the highly active viral Acr enzymes (16), suggesting that cellular Crn2 enzymes are not always “de-tuned” to act more slowly than their viral cousins. Furthermore, the existence of a fusion between Csx1 and Crn2 means that the relative concentrations of the two enzymes can not be varied in response to infection – they are of necessity fixed at 1:1. This is a distinct advantage when exploring the logic of a complex system as the relative concentrations of the effectors are defined with certainty.

These observations raise the intriguing question of how the two activities collaborate to provide appropriate levels of immunity. Our *in vitro* analyses clearly demonstrate that the ring nuclease activity limits, but does not abolish, the cA_4_-activated ribonuclease activity of Csx1 (Figure 4). One can rationalise this by assuming that the two enzymes compete for binding of cA_4_, and that once activated, the Csx1 enzyme can catalyse multiple rounds of RNA cleavage before the activator dissociates. Only then is there a chance to degrade it at the Crn2 active site. The balance between ribonuclease and ring nuclease enzyme activities will thus largely be determined by their relative affinities for cA_4_. Perhaps surprisingly, we observe that the Crn2 domain binds cA_4_ much more tightly than the CARF domain (apparent K_D_ ∼20 nM and > 1 µM, respectively). This suggests that the RNase activity of Csx1 will only be licensed once cA_4_ concentrations reach high µM levels in the cell. This is not necessarily a limitation, as the large degree of signal amplification generated by type III CRISPR systems can result in a 6 µM concentration of cA_4_ in cells detecting only 1 viral RNA (13). It should also be noted that a second cA_4_-activated defence enzyme is encoded by the CRISPR locus of *M. piezophila* – a predicted RelE enzyme that may be licensed by cA_4_ binding to cleave mRNA undergoing translation. Thus the Crn2 ring nuclease may need to function in the deactivation of two enzymes that could be toxic to the cell if not restrained. Overall, this study highlights the pressing need for prokaryotic cells to have a mechanism to remove cyclic nucleotide second messengers efficiently in circumstances where abortive immunity is not the desired outcome.

## METHODS

### Cloning and expression

For cloning, a synthetic gene (g-block) encoding the Csx1-Crn2 protein, codon optimised for *Eschericia coli*, was purchased from Integrated DNA Technologies (IDT), Coralville, USA, and cloned into the vector pV5HisTev (20) between the *Nco*I and *Bam*HI sites. Competent DH5α cells were transformed with the construct and sequence integrity confirmed by sequencing (Eurofins Genomics). A single conservative mutation (V177L) in a variable region of the protein was noted in the cloned gene but was not deemed problematical. The plasmid was transformed into *Escherichia coli* C43 (DE3) cells for protein expression. 2 L of LB culture was grown at 37 °C to an OD_600_ of 0.3 with shaking at 180 rpm. Protein expression was induced with 0.4 mM Isopropyl β-D-1-thiogalactopyranoside and cells were grown at 37 °C overnight before harvesting by centrifugation at 4000 rpm (Beckman Coulter Avanti JXN-26; JLA8.1 rotor) at 10 °C for 10 min.

### Protein purification

Expression and purification of *Sulfolobus islandicus* rod-shaped virus 1 gp29 (AcrIII-1) has been described previously (16). For Csx1-Crn2 purification, the cell pellet was resuspended in four volumes equivalent of lysis buffer containing 50 mM Tris-HCl 7.5, 0.5 M NaCl, 10 mM imidazole and 10 % glycerol supplemented with EDTA-free protease inhibitor tablets (Roche; 1 tablet per 100 ml buffer) and lysozyme (1 mg/ml). Cells were lysed by sonication (six times 1 min with 1 min rest intervals on ice) and the lysate was centrifuged at 40,000 rpm (70 Ti rotor) at 4 °C for 30 min. The lysate was filtered (0.45 µm) and then loaded onto a 5 ml HisTrap FF Crude column (GE Healthcare) equilibrated with wash buffer containing 50 mM Tris-HCl pH 7.5, 0.5 M NaCl, 30 mM imidazole and 10 % glycerol. Unbound protein was washed away with 20 column volumes (CV) of wash buffer prior to elution of his-tagged protein using a linear gradient (holding at 10 % for 3 CV, and 50 % for 3 CV) of elution buffer containing 50 mM Tris-HCl pH 7.5, 0.5 M NaCl, 0.5 M imidazole and 10 % glycerol. SDS-PAGE was carried out to identify fractions containing the protein of interest, and relevant fractions were pooled and concentrated using a 30 kDa molecular weight cut-off centrifugal concentrator (Merck). The his-tag was removed by incubating concentrated protein overnight with Tobacco Etch Virus (TEV) protease (1 mg per 10 mg protein) while dialyzing in buffer containing 50 mM Tris-HCl pH 7.5, 0.5 M NaCl, 30 mM imidazole and 10 % glycerol at room temperature. The protein with his-tag removed was filtered (0.45 µm) and isolated using a 5 ml HisTrapFF column, eluting the protein using 4 CV wash buffer. Cleaved protein was filtered (0.45 µm) and further purified by size-exclusion chromatography (Superdex S200 26/60; GE Healthcare) in buffer containing 20 mM Tris pH 7.5, 0.25 M NaCl using an isocratic gradient. After SDS-PAGE, fractions containing protein of interest were concentrated and protein was aliquoted and stored at −80 °C.

Variant enzymes were generated using the QuickChange Site-Directed Mutagenesis kit as per manufacturer’s instructions (Agilent technologies), verified by DNA sequencing and purified as for the wild-type protein.

### RNase assays

The RNA oligonucleotide A1 was purchased from IDT and RNA labelling was carried out as described previously (4). For RNA degradation assays, 50 nM radiolabelled RNA A1 was incubated with Csx1-Crn2 (5 or 10 µM dimer) with 100 µM of cyclic oligoadenylate (cA_3_, cA_4_, or cA_6_) in Reaction buffer (20 mM MES pH 6.5, 150 mM NaCl, 1 mM EDTA, 1 mM DTT and 0.06 U/µl SUPERase•In RNase Inhibitor) for 30 min (or other indicated timepoints), at 50 °C. Reactions were quenched by adding phenol-chloroform-isoamyl alcohol (Ambion) and vortexing. For single-turnover RNA cleavage kinetics experiments, a mix of 50 nM radiolabelled RNA A1 and 100 µM cA_4_ was prepared in pH 6.5 reaction buffer as above and pre-warmed at 50 °C for 5 min. Addition of pre-warmed enzyme (10 µM dimer Csx1-Crn2 or its variants) started the reaction. At indicated times, 10 µl of the reaction was taken and quenched by adding to phenol-chloroform and vortexing. Subsequently, the deproteinised RNA cleavage products were mixed (1:1) with 100% formamide xylene-cyanol loading dye and separated by denaturing polyacrylamide gel electrophoresis (PAGE; 20% acrylamide, 7M Urea, 1X TBE, 45 °C, 3000V) for 2 h. Following the electrophoresis, the gels were phosphorimaged at −70 °C overnight. For RNA kinetic analysis, substrate RNA and degradation products were quantified using the Bio-Formats plugin of ImageJ as distributed in the Fiji package and a background signal was subtracted as described previously (20). Fraction cleaved was calculated for every time point and fitted via GraphPad Prism 8 to an exponential graph Y=Y0 + (Plateau-Y0) *(1-exp(-K*x)) with Y0 fixed to 0 (One phase association). A1 oligo: 5’-AGGGUAUUAUUUGUUUGUUUCUUCUAAACUAUAAGCUAGUUCUGGAGA

### cA_4_ pre-incubation assay

Prior to the addition of RNA, 10 µM enzyme dimer WT or its variant H495A were mixed with 10, 50, or 100 µM cA_4_ in Reaction buffer and incubated for 10 min at 50 °C. To start the reaction, ∼50 nM radiolabelled RNA A1 was added and the reactions were incubated for a further 30 min at the same temperature before quenching and analysis as described above.

### cA_4_ degradation assays

Radiolabelled cyclic oligoadenylates were generated enzymatically using type III-D effector complex of *Sulfolobus solfataricus*, as described previously (14). For cA_4_ cleavage assay, ∼200 nM radiolabelled cA_4_ was incubated with either 1 or 5 μM dimer Csx1-Crn2 or its variants in Reaction buffer at 50 °C, over a set of time points as indicated in the relevant figure legends. As a positive control, cA_4_ and 1 µM AcrIII-1 SIRV1 gp29 were assayed for 1 min. At the desired time points, 10 µl of the reaction was removed and quenched by adding to phenol-chloroform and vortexing. Deproteinised products were then further isolated by chloroform-isoamyl alcohol extraction (Sigma-Aldrich) for thin-layer chromatography. As a negative control, cA_4_ was incubated to the end-point of each experiment.

### Multiple-turnover kinetics

For multiple turnover kinetics of cA_4_ cleavage, a reaction mix composed of 256 µM unlabelled cA_4_, ∼100 nM radiolabelled cA_4_ and varying concentration of enzyme dimer (160, 320, 480, 640, 1280, 2560 and 5120 nM) in Reaction buffer was prepared and incubated at 50 °C. Addition of pre-warmed enzyme to the reaction mix started the reaction. At certain time points, 10 µl of the reaction was removed, quenched, and extracted as described above. RNA cleavage was then visualised by denaturing PAGE and the fraction of RNA cut was quantified and initial rates were calculated. Initial rates were plotted against substrate concentration and fitted to a linear equation on GraphPad Prism 8 to obtain the multiple-turnover rate of cA_4_ cleavage by Csx1-Crn2.

### Thin-layer chromatography (TLC)

TLC was carried out as described previously (16). In brief, a glass chamber was pre-warmed for 1 hour at 38 °C and then left to humidify with 0.5 cm of TLC buffer (100 ml; 0.2 M ammonium bicarbonate pH 9.2, 70% ethanol, H_2_O) for a further 45 min. 1 µl of sample was spotted 1 cm from the bottom of a 20 cm x 20 cm silica gel on TLC Aluminium foil plate (Supelco Sigma-Aldrich) and the TLC plate was placed into the humidified glass chamber, sealed, and the buffer was allowed to migrate up the plate for ∼3 h at 35 °C (until the migration front reached ∼17 cm). The TLC plate was then air dried and phosphorimaged overnight.

### Electrophoretic mobility shift assay

For the binding assay, 10 nM radioactively-labelled cA_4_ was incubated with Csx1-Crn2 variants H495A, H129W and S452Y at varying concentrations in binding buffer containing 20 mM Tris–HCl pH 7.5, 150 mM NaCl and 2 mM MgCl_2_ supplemented with 2 μM Ultrapure Bovine Serum Albumin (Invitrogen) for 10 min at 25 °C. Following the incubation, samples were mixed with an equivalent volume of 20 % (v/v) glycerol and equal volumes were run on a native polyacrylamide gel (15 % acrylamide, 1 X TBE) at 250 V, 30 W and 28 °C for ∼3 h. Binding was visualized by phosphorimaging at −70 °C overnight.

## Author contributions

A.S., designed experiments, carried out experiments and analysed data in consultation with J.S.A and M.F.W. J.SA., helped A.S. design experiments, carried out experiments and formal analysis. S.G., cloned and purified proteins. M.F.W., conceptualised the project, obtained funding, carried out formal analysis and prepared the original draft of the manuscript. All authors contributed to writing, review and editing.

## References

1. Samai, P., Pyenson, N., Jiang, W., Goldberg, G.W., Hatoum-Aslan, A. and Marraffini, L.A. (2015) Co-transcriptional DNA and RNA Cleavage during Type III CRISPR-Cas Immunity. Cell, 161, 1164–1174.

2. Elmore, J.R., Sheppard, N.F., Ramia, N., Deighan, T., Li, H., Terns, R.M. and Terns, M.P. (2016) Bipartite recognition of target RNAs activates DNA cleavage by the Type III-B CRISPR-Cas system. Genes Dev, 30, 447–459.

3. Estrella, M.A., Kuo, F.T. and Bailey, S. (2016) RNA-activated DNA cleavage by the Type III-B CRISPR-Cas effector complex. Genes Dev, 30, 460–470.

4. Rouillon, C., Athukoralage, J.S., Graham, S., Grüschow, S. and White, M.F. (2018) Control of cyclic oligoadenylate synthesis in a type III CRISPR system. eLife, 7, e36734.

5. Niewoehner, O., Garcia-Doval, C., Rostol, J.T., Berk, C., Schwede, F., Bigler, L., Hall, J., Marraffini, L.A. and Jinek, M. (2017) Type III CRISPR-Cas systems produce cyclic oligoadenylate second messengers. Nature, 548, 543–548.

6. Kazlauskiene, M., Kostiuk, G., Venclovas, C., Tamulaitis, G. and Siksnys, V. (2017) A cyclic oligonucleotide signaling pathway in type III CRISPR-Cas systems. Science, 357, 605–609.

7. Nasef, M., Muffly, M.C., Beckman, A.B., Rowe, S.J., Walker, F.C., Hatoum-Aslan, A. and Dunkle, J.A. (2019) Regulation of cyclic oligoadenylate synthesis by the S. epidermidis Cas10-Csm complex. RNA, 25, 948–962.

8. Foster, K., Kalter, J., Woodside, W., Terns, R.M. and Terns, M.P. (2019) The ribonuclease activity of Csm6 is required for anti-plasmid immunity by Type III-A CRISPR-Cas systems. RNA Biol, 16, 449–460.

9. Jia, N., Jones, R., Yang, G., Ouerfelli, O. and Patel, D.J. (2019) CRISPR-Cas III-A Csm6 CARF Domain Is a Ring Nuclease Triggering Stepwise cA4 Cleavage with ApA>p Formation Terminating RNase Activity. Mol. Cell, 75, 933–943.

10. Molina, R., Stella, S., Feng, M., Sofos, N., Jauniskis, V., Pozdnyakova, I., Lopez-Mendez, B., She, Q. and Montoya, G. (2019) Structure of Csx1-cOA4 complex reveals the basis of RNA decay in Type III-B CRISPR-Cas. Nature communications, 10, 4302.

11. McMahon, S.A., Zhu, W., Rambo, R., Graham, S., White, M.F. and T.M., G. (2020) Structure and mechanism of a Type III CRISPR defence DNA nuclease activated by cyclic oligoadenylate. Nature Commun., 11, 500.

12. Rostol, J.T. and Marraffini, L.A. (2019) Non-specific degradation of transcripts promotes plasmid clearance during type III-A CRISPR-Cas immunity. Nat Microbiol, 4, 656–662.

13. Athukoralage, J.S., Graham, S., Rouillon, C., Grüschow, S., Czekster, C.M. and White, M.F. (2020) The dynamic interplay of host and viral enzymes in type III CRISPR-mediated cyclic nucleotide signalling. BioRxiv, 2020.02.12.946046.

14. Athukoralage, J.S., Rouillon, C., Graham, S., Grüschow, S. and White, M.F. (2018) Ring nucleases deactivate Type III CRISPR ribonucleases by degrading cyclic oligoadenylate. Nature, 562, 277–280.

15. Athukoralage, J.S., Graham, S., Grüschow, S., Rouillon, C. and White, M.F. (2019) A type III CRISPR ancillary ribonuclease degrades its cyclic oligoadenylate activator. J. Mol. Biol., 431, 2894–2899.

16. Athukoralage, J.S., McMahon, S.A., Zhang, C., Gruschow, S., Graham, S., Krupovic, M., Whitaker, R.J., Gloster, T.M. and White, M.F. (2020) An anti-CRISPR viral ring nuclease subverts type III CRISPR immunity. Nature, 577, 572–575.

17. Waterhouse, A., Bertoni, M., Bienert, S., Studer, G., Tauriello, G., Gumienny, R., Heer, F.T., de Beer, T.A.P., Rempfer, C., Bordoli, L. et al. (2018) SWISS-MODEL: homology modelling of protein structures and complexes. Nucl. Acids Res., 46, W296–W303.

18. Lapinaite, A., Doudna, J.A. and Cate, J.H.D. (2018) Programmable RNA recognition using a CRISPR-associated Argonaute. Proc. Natl. Acad. Sci. USA, 115, 3368–3373.

19. Griffin, M.A., Davis, J.H. and Strobel, S.A. (2013) Bacterial toxin RelE: a highly efficient ribonuclease with exquisite substrate specificity using atypical catalytic residues. Biochemistry, 52, 8633–8642.

20. Rouillon, C., Athukoralage, J.S., Graham, S., Grüschow, S. and White, M.F. (2019) Investigation of the cyclic oligoadenylate signalling pathway of type III CRISPR systems. Methods Enzymol, 616, 191–218.

21. Grüschow, S., Athukoralage, J.S., Graham, S., Hoogeboom, T. and White, M.F. (2019) Cyclic oligoadenylate signalling mediates Mycobacterium tuberculosis CRISPR defence. Nucl. Acids Res., 47, 9259–9270.

22. Yan, X., Guo, W. and Yuan, Y.A. (2015) Crystal structures of CRISPR-associated Csx3 reveal a manganese-dependent deadenylation exoribonuclease. RNA Biol, 12, 749–760.

23. Han, W., Pan, S., Lopez-Mendez, B., Montoya, G. and She, Q. (2017) Allosteric regulation of Csx1, a type IIIB-associated CARF domain ribonuclease by RNAs carrying a tetraadenylate tail. Nucl. Acids Res., 45, 10740–10750.

24. Lau, R.K., Ye, Q., Birkholz, E.A., Berg, K.R., Patel, L., Mathews, I.T., Watrous, J.D., Ego, K., Whiteley, A.T., Lowey, B. et al. (2020) Structure and Mechanism of a Cyclic Trinucleotide-Activated Bacterial Endonuclease Mediating Bacteriophage Immunity. Mol. Cell, 77, 723–733.

25. Ye, Q., Lau, R.K., Mathews, I.T., Birkholz, E.A., Watrous, J.D., Azimi, C.S., Pogliano, J., Jain, M. and Corbett, K.D. (2020) HORMA Domain Proteins and a Trip13-like ATPase Regulate Bacterial cGAS-like Enzymes to Mediate Bacteriophage Immunity. Mol. Cell, 77, 709–722 e707.

26. Cohen, D., Melamed, S., Millman, A., Shulman, G., Oppenheimer-Shaanan, Y., Kacen, A., Doron, S., Amitai, G. and Sorek, R. (2019) Cyclic GMP-AMP signalling protects bacteria against viral infection. Nature, 574, 691–695.

27. Whiteley, A.T., Eaglesham, J.B., de Oliveira Mann, C.C., Morehouse, B.R., Lowey, B., Nieminen, E.A., Danilchanka, O., King, D.S., Lee, A.S.Y., Mekalanos, J.J. et al. (2019) Bacterial cGAS-like enzymes synthesize diverse nucleotide signals. Nature, 567, 194–199.

